# Rapid and accurate discrimination of *Mycobacterium abscessus* subspecies based on matrix-assisted laser desorption ionization-time of flight spectrum and machine learning algorithms

**DOI:** 10.1101/2022.09.06.506122

**Authors:** Hsin-Yao Wang, Chi-Heng Kuo, Chia-Ru Chung, Ting-Wei Lin, Jia-Ruei Yu, Jang-Jih Lu, Ting-Shu Wu

**Author notes:** These authors contribute equally to the work (Hsin-Yao Wang, Chi-Heng Kuo). To whom correspondence should be addressed: JJ Lu, TS Wu, **CORRESPONDENCE**: Hsin-Yao Wang, MD, Address: No. 5, Fuxing St., Guishan Dist., Taoyuan City 333, Taiwan.

## Abstract

**Background:** *Mycobacterium abscessus* complex (MABC) has been reported to cause considerable complicated infections. Subspecies identification of MABC is crucial for adequate treatment due to different antimicrobial resistance property amid the subspecies. However, long incubation days is needed for the traditional antibiotic susceptibility testing (AST) method. Effective antibiotics administration often delayed considerably and caused unfavorable outcomes. Thus, we proposed a novel and accurate method to identify subspecies and its potential antibiotics resistance, to guide clinical treatment within hours.

**Methods:** Subspecies of the MABC isolates were determined by *secA1, rpoB*, and *hsp65*. AST was tested by using microdilution method, as well as sequencing of *erm*(41) and *rrl* genes. MALDI-TOF mass spectrometry (MS) spectra were analyzed. The informative peaks on MS spectra were detected by random forest (RF) importance. Machine learning (ML) algorithms were used to build models for classifying MABC subspecies based on MALDI-TOF spectrum. The models were developed and validated by nested five-fold cross-validation to avoid over-fitting.

**Results:** In total, 102 MABC isolates (52 subspecies *abscessus* and 50 subspecies *massiliense*) were analyzed. Top informative peaks including *m/z* 6715, 4739, 2805, etc. were identified. RF model attained AUROC of 0.9166 (95% CI: 0.9072-0.9196) and outperformed other algorithms in discriminating subspecies *abscessus* from *massiliense*.

**Conclusion:** We developed a MALDI-TOF based ML model for rapid and accurate MABC subspecies identification. The novel diagnostic tool would guide a more accurate and timely MABC subspecies-specific treatment.

## Introduction

Nontuberculous mycobacterial (NTM) infection has become an emerging crisis worldwide due to its rising mortality rates. As most countries, *M. abscessus* complex (MABC) is one of the predominant NTM species in Taiwan.^1^ Specifically, MABC has been reported to cause 21.4% NTM infection.^2^ In European countries, MABC is the most common pathogen causing lung infection among cystic fibrosis patients, and the incidence is increasing.^3^ While it is well-acknowledged that treating MABC is much more challenging than other NTMs, rapid and accurate identification of MABC subspecies as well as the antibiotic susceptibility testing (AST) have raised considerable interest. However, the clinical need has not been well met. To diagnose MABC, at least two sputum samples that collected on different days for culture still serve as the microbiological standard of diagnosis in guidelines.^4^ Subspecies identification of MABC and AST are of utmost importance due to different response to macrolides amid the subspecies, leading to different treatment strategies.^5,6^ Traditional AST required at least 10 – 14 incubation days to determine whether drug-resistance developed, which postpones the treatment. The time delay between clinical visit and adequate treatment would worsen the infection and lead to unfavorable outcomes. There is a need to develop a more rapid approach to identify MABC subspecies and resistance to antibiotics. Thus, a novel and time-saving method would have a huge impact on current practices of NTM infection management.

MABC is a group of rapidly-growing species, remaining the most difficult one to deal with due to its inducible macrolide resistance.^6^ MABC can be divided into three main subspecies– *M. abscessus* subsp. *abscessus, M. abscessus* subsp. *massiliense* and *M. abscessus* subsp. *bolletii*.^7,8^ Functional *erm*(41) gene has been proven to activate macrolide resistance in subsp. *abscessus* and subsp. *bolletii*., leading to drug resistance after incubation to 10 – 14 days.^9^ In contrast, subsp. *massiliense* possesses dysfunctional *erm*(41) gene preventing from macrolide resistance.^6,10^ With the need to identify MABC, matrix-assisted laser desorption ionization-time of flight mass spectrometry (MALDI-TOF MS)^11^ is a more rapid, inexpensive approach with high accuracy for differentiation of subspecies compared with conventional approach such as phenotypic methods and *rpoB* gene–based sequencing.^12,13^ MALDI-TOF MS spectrum can be easily used to identify different subspecies among MABC.^13^ However, traditional interpretation of complex MALDI-TOF MS spectrum data solely by investigators is operator-dependent and consequently not robust.

Machine learning (ML) technology has been introduced to save considerable time in analyzing complex data, without human error.^14,15^ The ML-based methods have been reported for rapid and accurate detection of resistant bacteria (e.g., *E. coli*,^16^ *S. aureus*,^17^ *E. faecium*^18^) from MALDI-TOF MS spectrum. Additional, subspecies identification can be achievable by applying ML in analyzing MALDI-TOF MS spectrum.^19,20^ ML techniques, including support vector machines (SVM), random forests (RFs), logistic regression (LR), decision tree (DT), and k-nearest neighbor (KNN) have been widely implemented to analyze microbiological features and construct model of classification.^11^ Thus, harnessing ML in interpreting MALDI-TOF MS spectrum is a reasonable approach to provide rapid and accurate identification of MABC subspecies.

In present study, we proposed a novel approach to discriminate MABC subspecies by integrating MALDI-TOF and ML method. Based on the molecular typing and antibiotic susceptibility tests of MABC, we demonstrated the novel approach provides rapid subspecies identification for MABC, which can lead to a more adequate management of MABC infectious disease.

## Methods

### Study protocol

Overall research scheme was presented as **Figure 1**. The new approach could save plenty of time in MABC subspecies identification and the corresponding AST. The workflow of developing ML models was illustrated in **Figure 2**, mainly included four parts: 1) sample preparation, 2) MS spectra and data processing, 3) model training and validation, 4) model testing and performance valuation. Isolates were collected at Chang Gung Memorial Hospital (CGMH) Linkou branch, Taiwan, from August 1^st^ 2015 to March 31^st^, 2018. Specimens were all collected from the sputum. After specimen collection and laboratory preparation, MALDI-TOF MS spectra were then obtained and feature selection was processed by ML methods for construction of predictive models. Robust and unbiased model validation was then done using nested five-fold cross-validation approach. Finally, after the classification models were created, we further calculated the model performance according to various ML algorithms.

**Figure 1.**
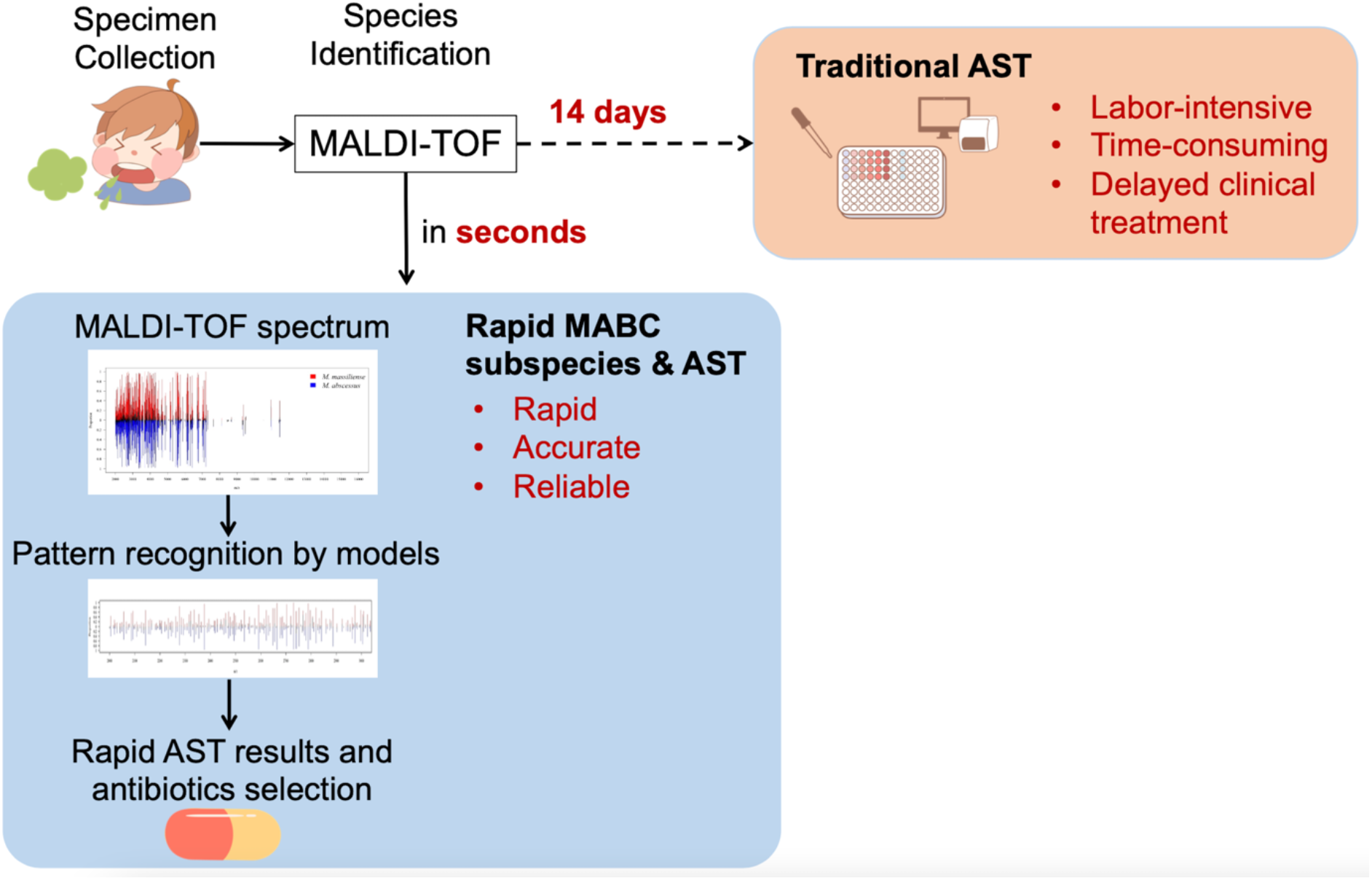
Comparison of the workflows. Antimicrobial susceptibility testing (AST) for MABC typically costs 14 days. By contrast, in the current study we propose a new MALDI-TOF based workflow for rapid MABC subspecies and AST identification in seconds. Based on the rapid and accurate MABC subspecies identification, an early and precise management for MABC makes possible.

**Figure 2.**
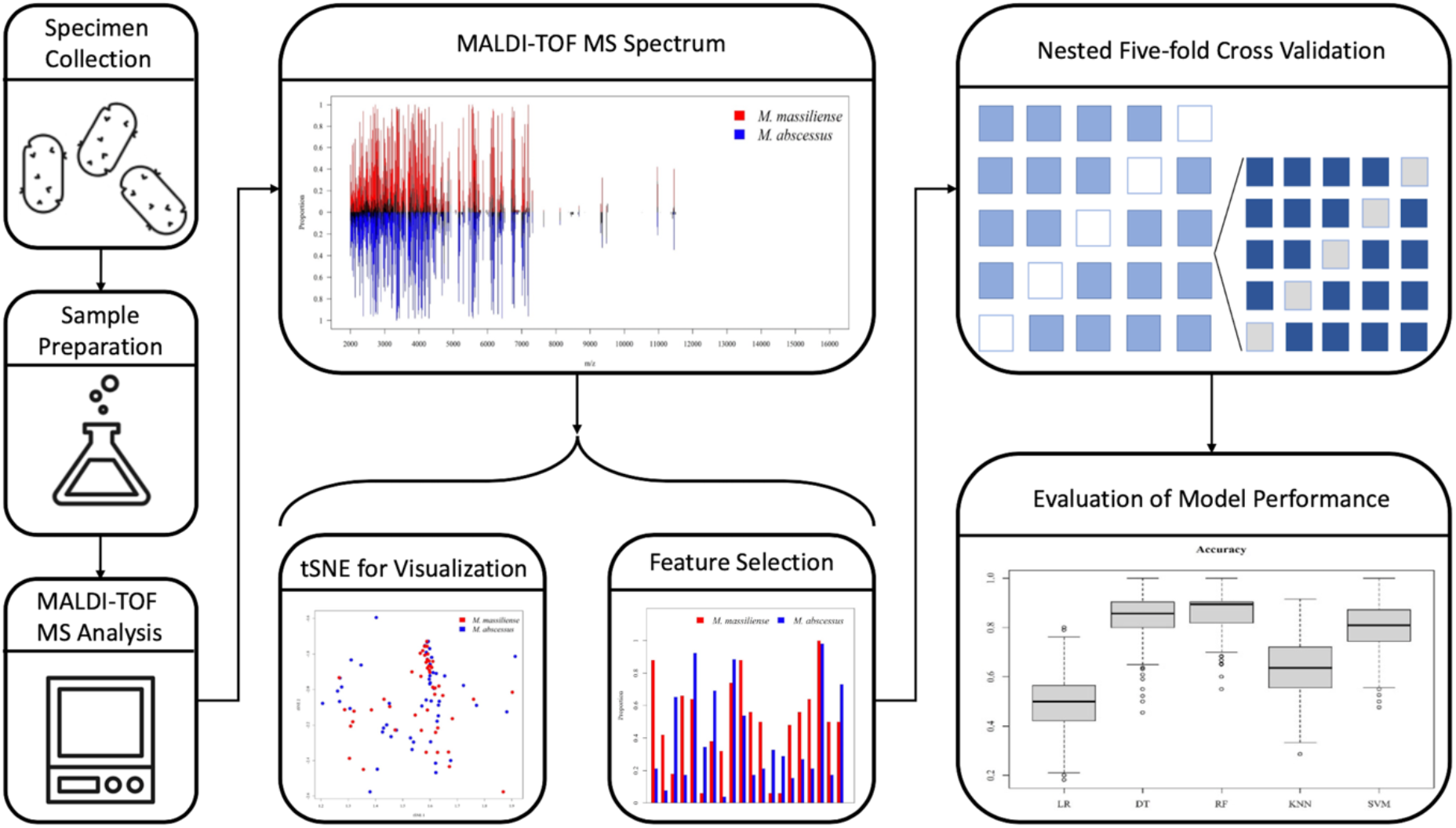
Scheme of the workflow. MABC isolates are analyzed by MALDI-TOF to retrieve MS spectra as the input. The subspecies classification for the MABC isolates are labeled according to DNA sequencing of the marker genes (i.e., *secA1, rpoB*, and *hsp65*). The MS spectra are analyzed and visualized prior to modeling. Informative features are also illustrated amid the subspecies. We train and validate the models by using nested cross-validation to avoid over-fitting. Performance metrics of different algorithms are calculated and compared.

### Sample preparation

Samples processing and preparation followed the laboratory protocol in CGMH hospital. Sputum samples were collected from the patients as daily routine. MABC isolates were stored at −70°C until analysis. One to three inoculation loops (10 μl loop) of bacterial stock were transferred into 300 μl ddH_2_O. The solution was then heated at 95°C for 30 minutes for inactivation. After heating, the solution was centrifuged at 13,000 rpm for two minutes. Repetitive washing (for three times) by dissolved in ddH_2_O and ethanol followed by centrifugation (13,000 rpm for two minutes) was conducted. After supernatant were completely removed, drying procedure was executed under room temperature for two minutes. Enhanced bacteria lysis was performed by adding 0.5 mm silica beads in 20 μl acetonitrile and maximally vortexed for 1 min. Twenty μl formic acid was added and then mixed well by vortex for five seconds, centrifugation at 13,000 rpm for two minutes was done again. One μl supernatant was loaded onto MALDI steel plate and dried in room temperature. One μl matrix solution (50% acetonitrile containing 1%α-cyano-4-hydroxycinnamic acid and 2.5% trifluoroacetic acid) was applied onto the plate, followed by dried in room temperature and analysis by MALDI-TOF MS.

### Measurement and Preprocessing of MALDI-TOF spectra

Microflex LT MS (Bruker Daltonics, Germany) was used to generate MALDI-TOF spectra for the isolates. Spectrum of mass-to-charge ratio (m/z) from 2,000 to 20,000 was collected. MS spectra was processed by using Flexanalysis 3.4. (Bruker Daltonics). MABC was identified according to Biotyper 3.1 (Bruker Daltonics). Default settings of manufacturer instruction were followed. The spectra were further processed by a modified bin method.^20^ In the processing step, features were extracted from the complex spectral data and formed well-defined structured data as the input features to ML models.

### Determination of MABC subspecies and AST

DNA sequences of *secA1, rpoB*, and *hsp65* were used as the markers for determination of MABC subspecies (i.e., subsp. *abscessus*, subsp. *massiliense*, and subsp. *bolletii*).^21^ Genes *erm*(41) and *rrl* were sequenced based on the method detailed in the previous study.^22^ AST of 11 antimicrobial agents (ie. amikacin, cefoxitin, ciprofloxacin, clarithromycin, doxycycline, imipenem, linezolid, minocycline, moxifloxacin, trimethoprim/sulfamethoxazole and tigecycline) were analyzed by broth microdilution (BMD). Specifically, the BMD was performed by using Sensititre RAPMYCOI MIC plates (Thermo Fisher). The reading time points for the BMD results included early reading time (3^rd^ to 5^th^ day) and late reading time (14^th^ day, specific for clarithromycin) to detect inducible macrolide resistance. Interpretation of AST for the 10 antimicrobial agents (except tigecycline) were based on CLSI M24 guideline.^23^

### Visual illustration of MS spectrum data onto a 2-dimensional plot

t-distributed stochastic neighbor embedding (tSNE) was designed to analyze multi-dimensional data to find clusters visually in a two-dimensional compressing data, by evaluating inter-cluster distance. tSNE is widely used to reduce the complex dimensionality retrieved from MALDI-TOF MS data.^24,25,26^ Initially, we analyzed our MS spectra, using the R-package “Rtsne” on R software in our research. We eventually produced a tSNE plot to visualize our analytic results.

### Feature selection and predictive model construction by ML method

Specific peaks showing distinct proportion between two subspecies, were selected as characteristic features. The difference reflected great potential to discriminate between unidentical subspecies. Feature selection was done before model construction, to include the crucial information to train models. Logistic regression (LR), decision tree (DT), random forest (RF), k-nearest neighbors (KNN), and support vector machine (SVM) were used to train and build the models. Random forest classifier was executed by ‘‘randomForest” library; SVM method was processed by “e1071” library; and “class” library was used for KNN algorithm; all of those packages were implemented with R software. The nested five-fold cross-validation process was then applied to validate the model.

### Model performance evaluation

Area under the receiver operating characteristic curve (AUROC) was calculated to evaluate the model performance. AUC of 0.5 represents random guess, i.e., tossing coins, thus AUROC of optimal models should be greater than 0.5 and as closer to 1. Youden’s J statistic was also introduced to generate sensitivity, specificity, and accuracy for the models.

## Results

### Bacterial isolates and MALDI-TOF MS spectra

In total, 102 MABC isolates were collected (**Table 1**), including 52 isolates classified as *M. abscessus* subsp. *abscessus* and other 50 isolates as *M. abscessus* subsp. *massiliense*. Regarding MALDI-TOF MS spectra, **Figure 3A** shows overall peaks distribution of the two subspecies over m/z 2,000 to 16,000. More detailed views on peaks distribution and difference of occurring frequency over m/z 2,000 to 6,000 were illustrated on **Figure 3B**, and m/z 6,000 to 10,000 in **Figure 3C**, respectively. Based on the information of peaks intensity, isolates of both subspecies were illustrated on a two-dimensional plot by tSNE method (**Figure 4**). As shown in the figure, data points of the two subspecies scattered loosely. Preliminarily, the two subspecies could not be discriminated clearly from each other, indicating classification by using higher dimensional information is needed.

**Figure 3.**
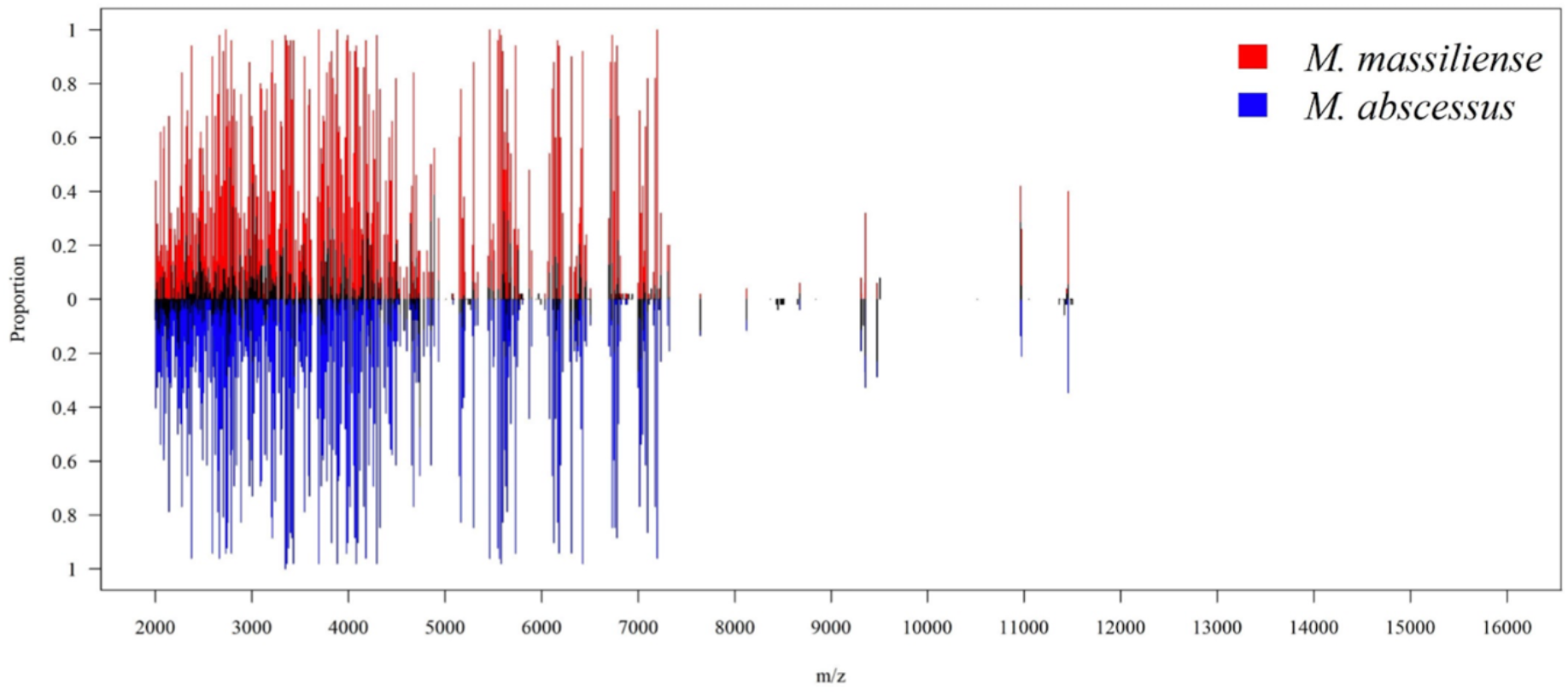
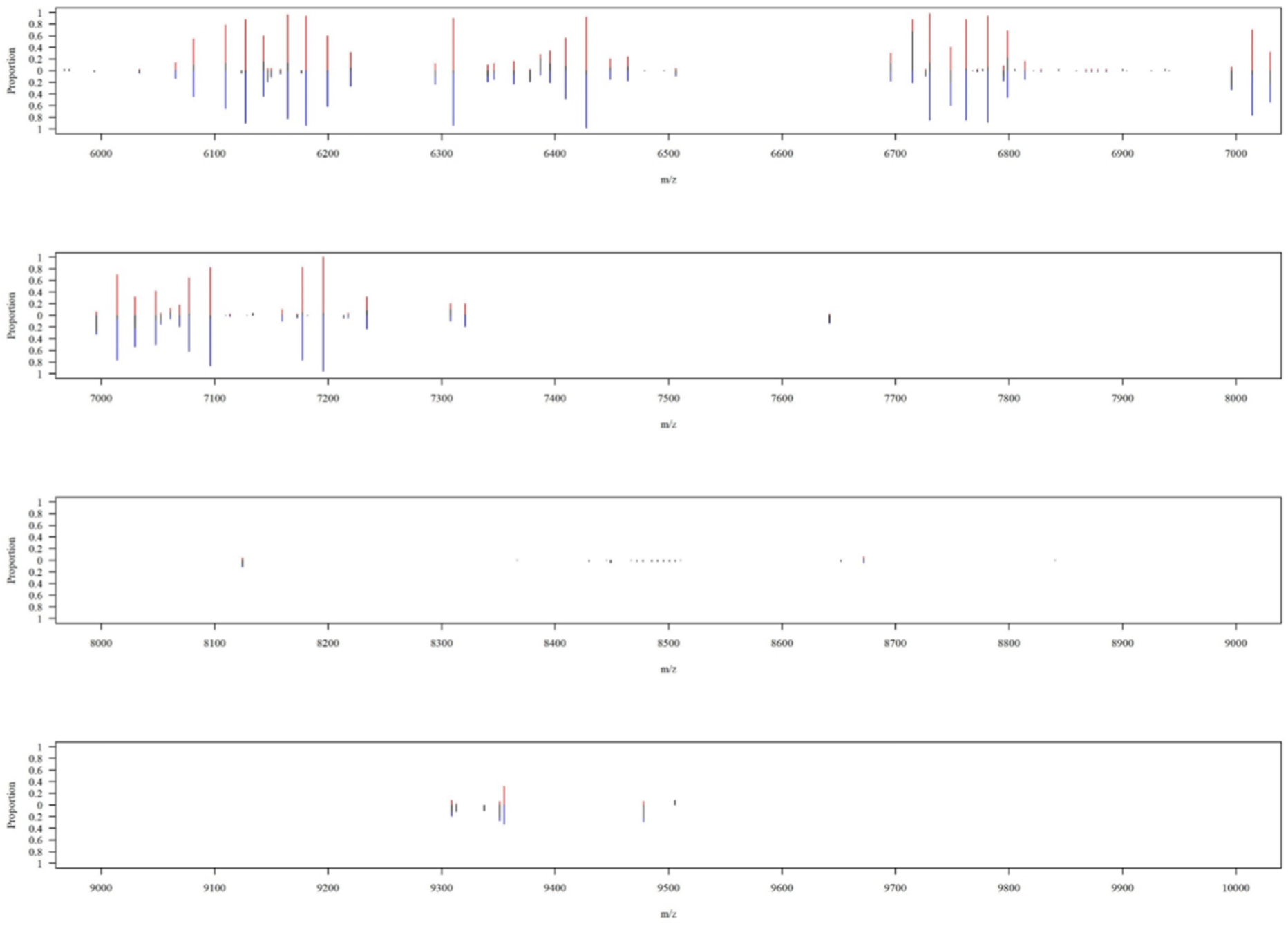
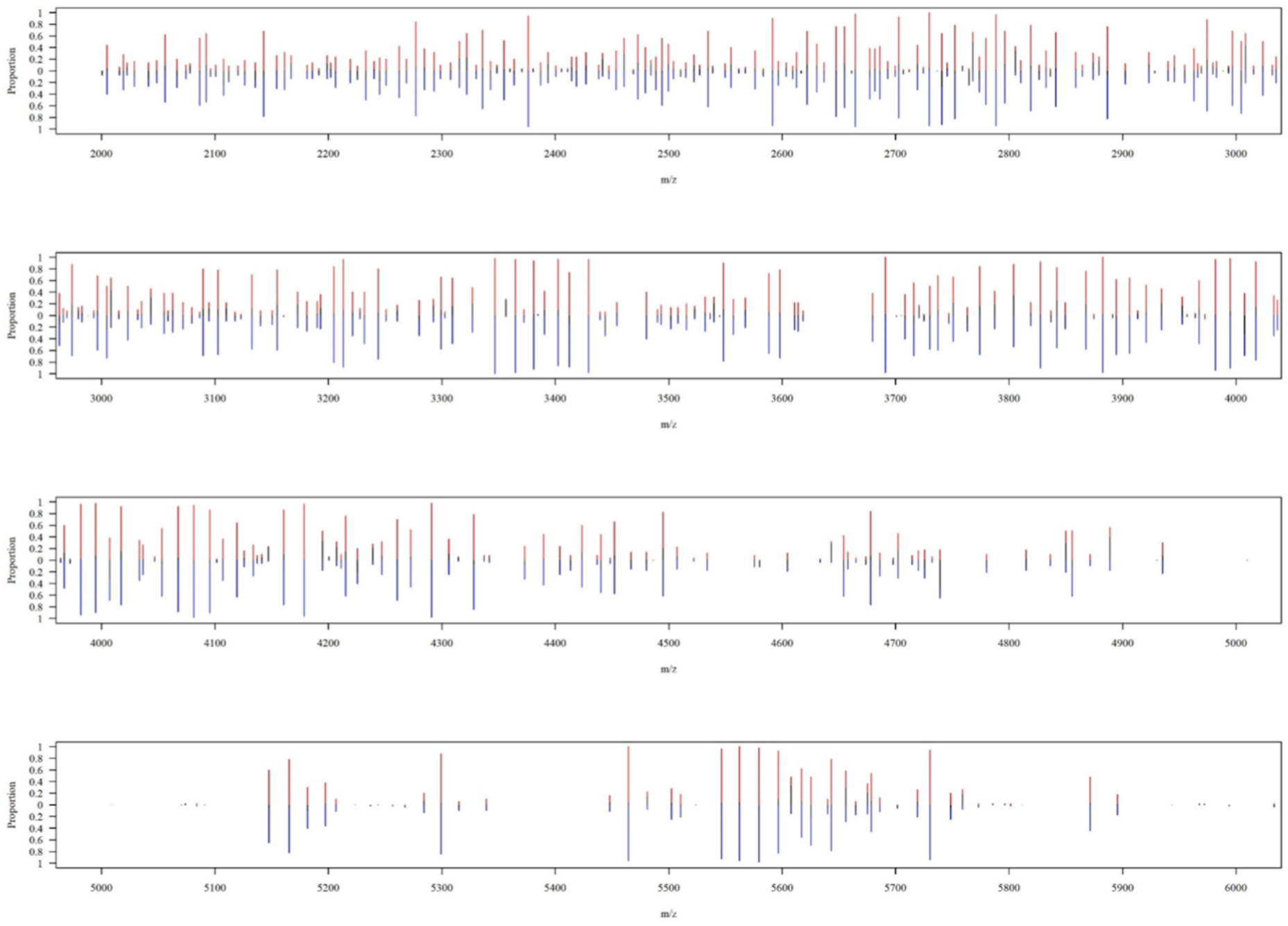
**(A).** Peaks distribution on MALDI-TOF spectrum of two subspecies of MABC over the range of m/z 2,000 to 16,000. Informative peaks are predominantly found over of 2,000 to 10,000. **(B).** A zoomed-in detailed view over the range of *m/z* 2,000 to 6,000. **(C).** A zoomed-in view over the range of m/z of 6,000 to 10,000. Peaks are mostly found at the range of *m/z* 2,000 to 6,000. MABC: *Mycobacterium abscessus* complex.

**Figure 4.**
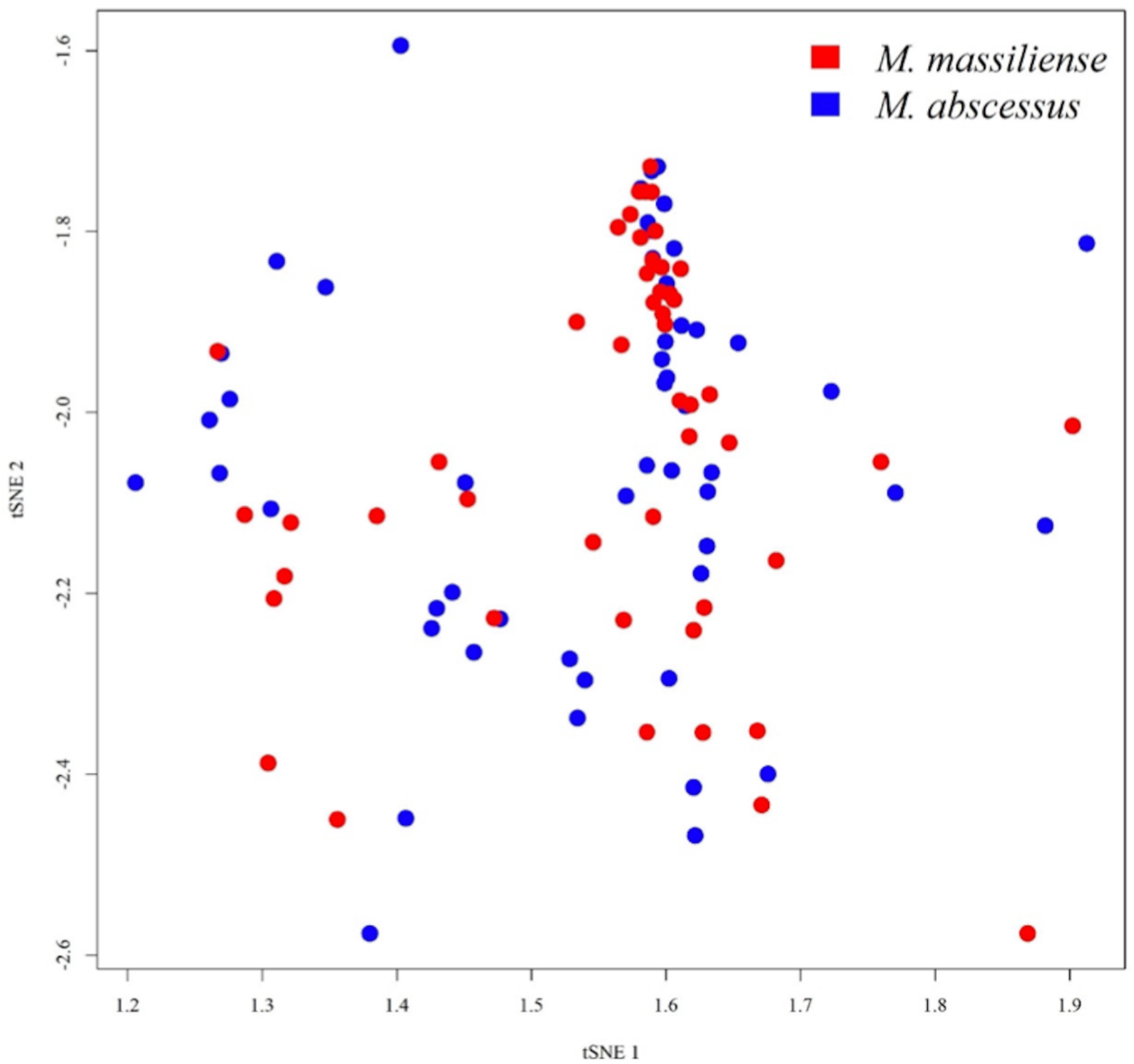
Visual illustration of MS spectrum data onto a two-dimensional plot. MALDI-TOF MS spectra of the two different MABC subspecies are analyzed and depict onto a two-dimensional plot. No obvious clustering or patterns can be observed based on the distribution of all the data points.

**Table 1.** Isolates number for *Mycobacterium abscessus* subsp. *abscessus* and *Mycobacterium abscessus* subsp. *massiliense*.

|  | *M. abscessus* | *M. massiliense* | Total |
| --- | --- | --- | --- |
| Number of isolates | 52 | 50 | 102 |
| Proportion | 51% | 49% |  |

### Performance of constructed models

Predictive performance of the ML models was shown in **Table 2**. In addition, box plots of the predictive performance metrics were presented in **Figure 5A**; we also drawn the ROC curves of predictive models, as shown in **Figure 5B**. LR attained sensitivity, specificity, accuracy, and AUROC as 0.4966, 0.4862, 0.4911, and 0.5729. This level of performance was not acceptable as a prediction tool. Similarly, KNN had better performance of sensitivity, specificity, accuracy, AUROC of 0.6353, 0.6404, 0.6380, 0.6849, but still not accurate enough to guide clinical decisions. DT, RF, and SVM had all the metrics above 0.80. Specifically for RF model, the sensitivity, specificity, accuracy, AUROC were 0.8647, 0.8711, 0.8695, 0.9166.

**Figure 5.**
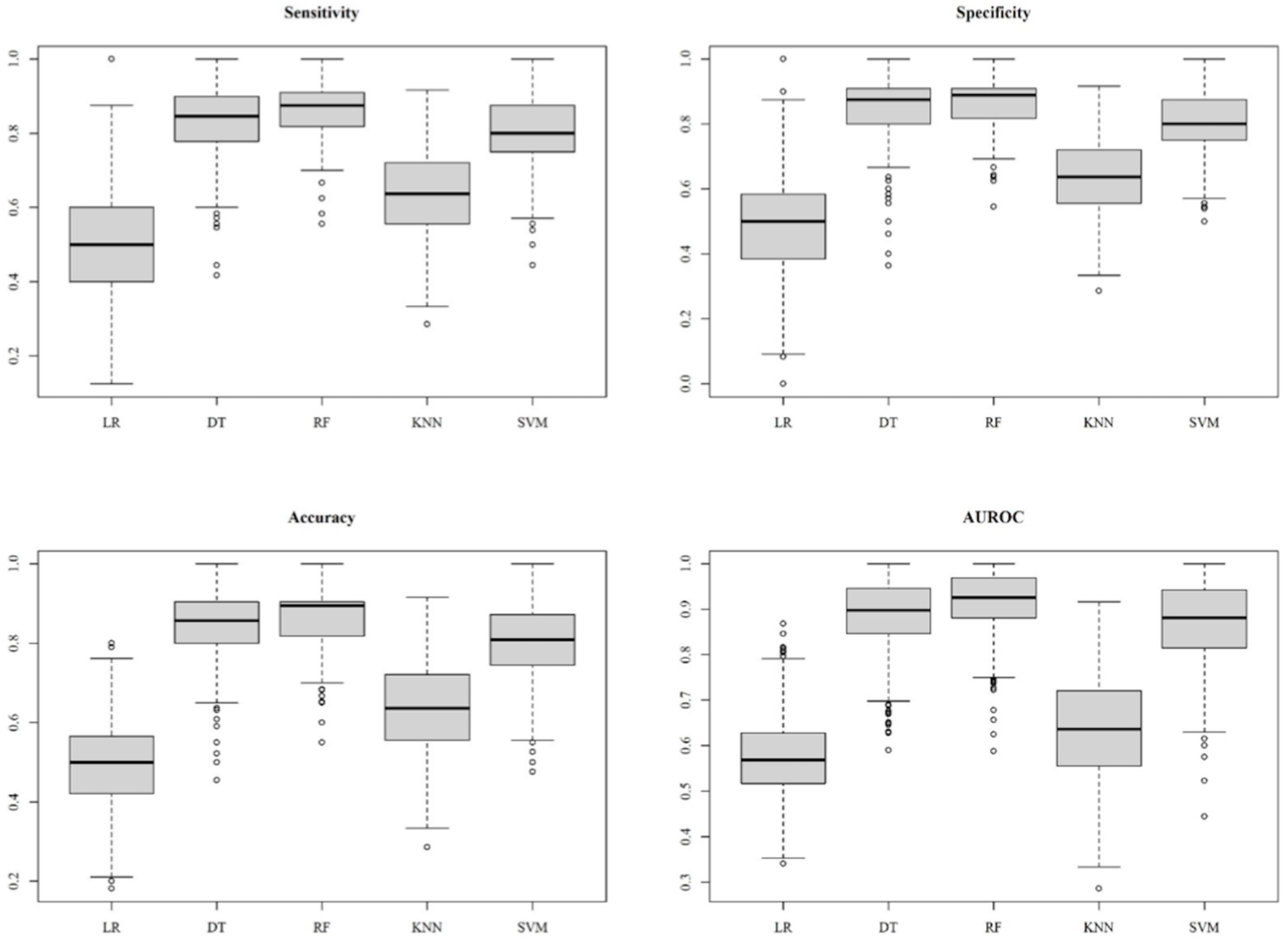
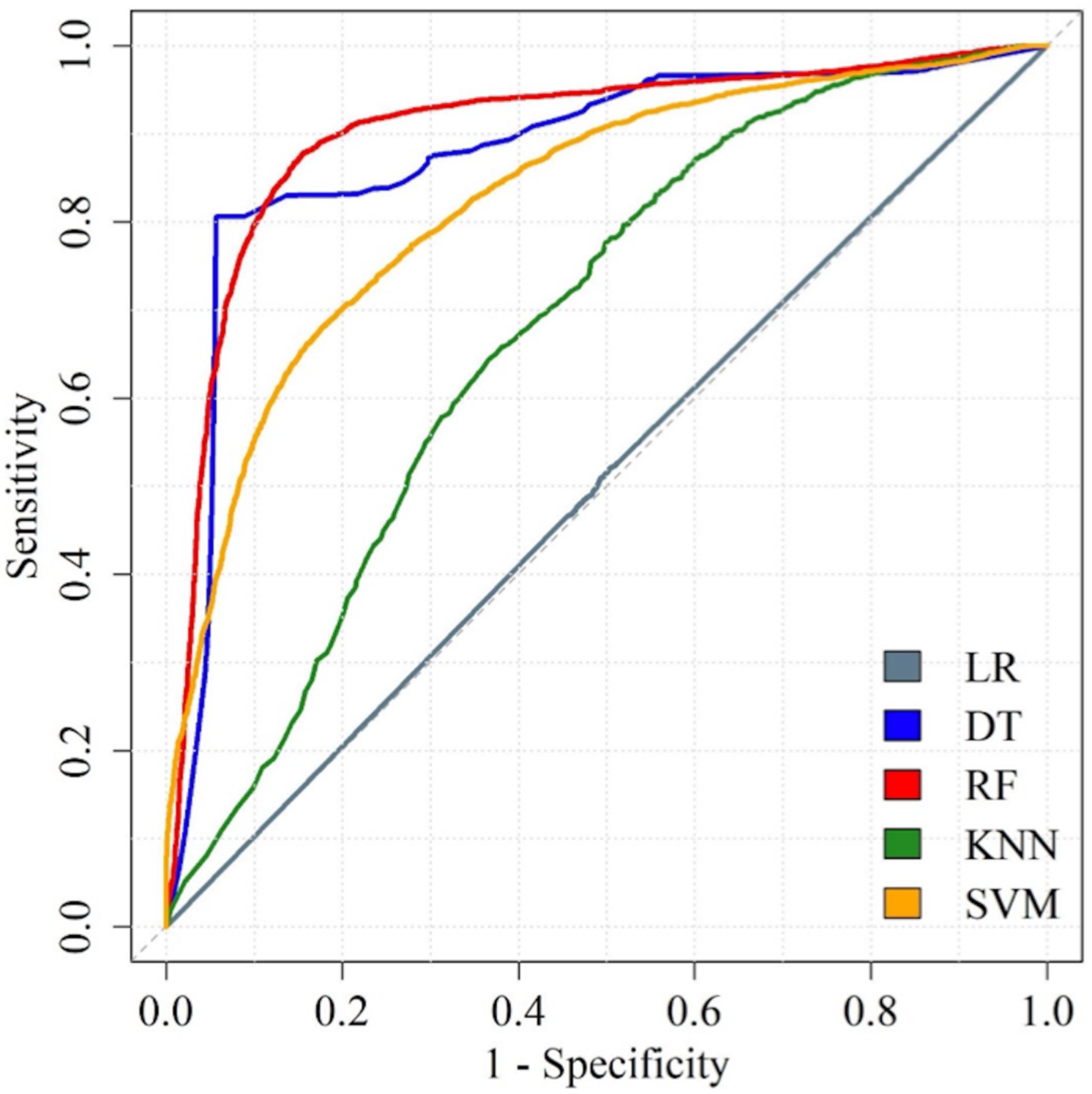
**(A).** Sensitivity, specificity, accuracy, and area under the receiver operating characteristic curve (AUROC) for the models with a various of algorithms. Amid the algorithms, RF and DT outperform other algorithms. **(B).** ROC curves of the five predictive models. LR: logistic regression; DT: decision tree; RF: random forest; KNN: k-nearest neighbor; SVM: support vector machine.

**Table 2.**
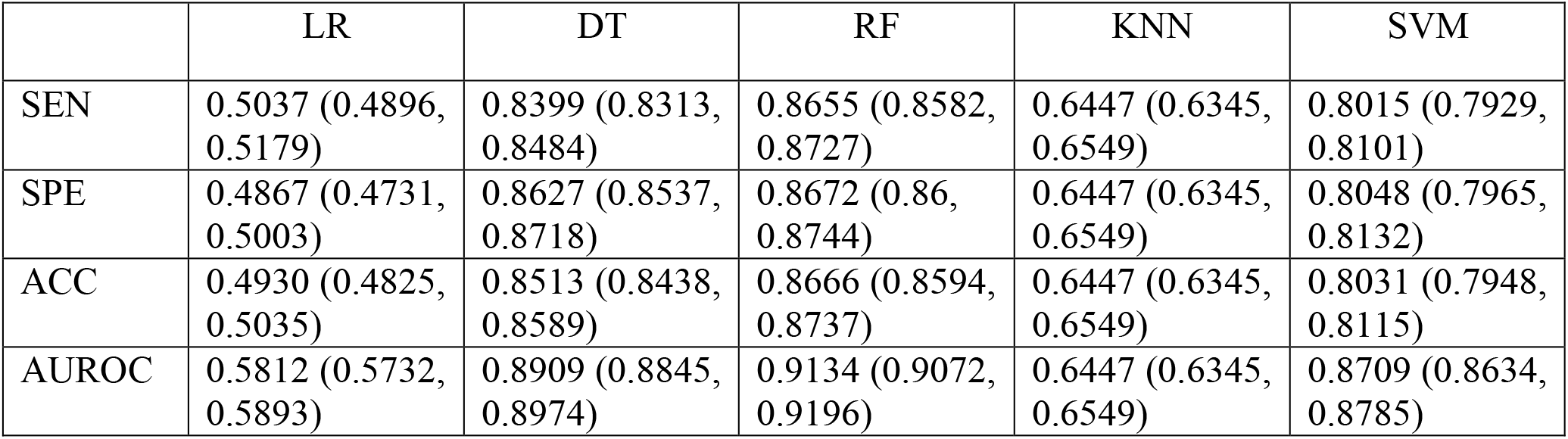
Diagnostic performance of the models. The metrics are expressed by the means and 95% confidence interval. SEN: sensitivity; SPE: specificity; ACC: accuracy; AUROC: receiver operating characteristic curve; LR: logistic regression; DT: decision tree; RF: random forest; KNN: k-nearest neighbor; SVM: support vector machine.

### Discriminative peaks

We found 20 informative peaks by RF importance method **(Figure 6A)**. As the Figure 6A showed, we ranked all 20 peaks in the order of Gini index. We further analyzed the occurring frequency of the peaks in *M. abscessus* subsp. *abscessus* and *M. abscessus* subsp. *massiliense*, as demonstrated in **Figure 6B**. Several peaks served as strong indicators. For instance, peaks of *m/z* 6,715, 2,805, 2,768, 4,643, 3,804, 4,889, 4850, 5,608, 2,460, 3,008, 4,195 predominantly suggested subsp. *massiliense* due to its high proportion in subsp. *massiliense* but low proportion in subsp. *abscessus*. In contrast, peaks of *m/z* 4,739, 2,741, 3,444, 4,007, 3,412, 6,996, 9,478, 3,005 implied subsp. *abscessus*.

**Figure 6.**
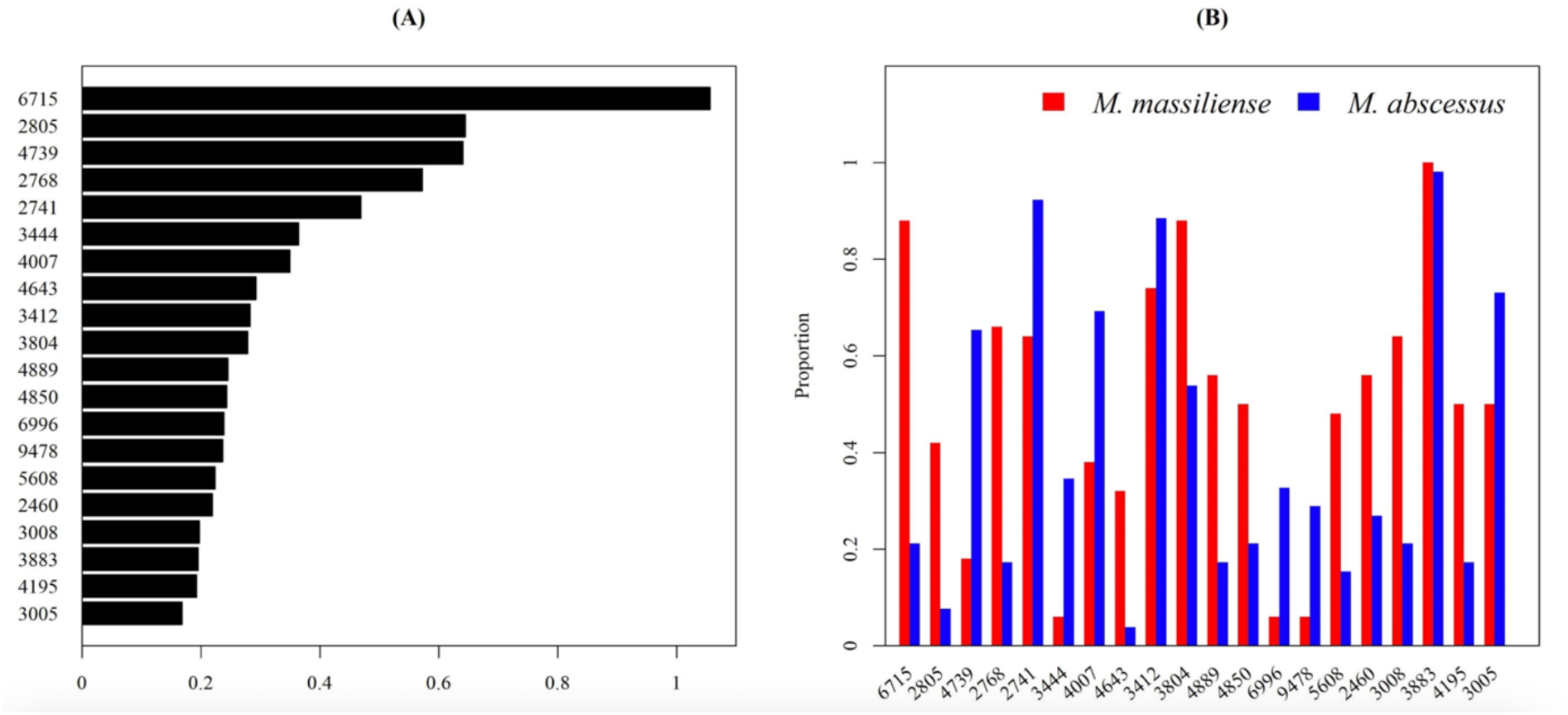
Informative peaks. (A) Informative peaks are sorted by RF importance. (B) The occurring frequency of the informative peaks amid the two MABC subspecies.

## Discussion

In this study, we offered a novel approach to differentiate MABC subspecies with drug-resistance (i.e., subsp. *abscessus*) from subspecies with drug-susceptibility (i.e., subsp*. massiliense*). Our study demonstrated ML algorithms can effectively interpret the implicit peaks pattern on a MALDI-TOF MS spectrum, and provide identification for MABC subspecies within seconds. The rapid and accurate subspecies classification would guide a more appropriate antibiotics administration for treating MABC.

Genetic test is typically the testing method that is used to identify MABC subspecies and antibiotic resistance. Previous studies showed that genetic changes by sequence analysis of *rpoB, hsp65, secA1*, 23S *rrl* and *erm*(41) genes accomplish classifying subspecies and specific antibiotics resistance accurately.^27,28,29^ Marras et al.^30^ reported that a real-time multiplex PCR assay approach using molecular beacons could identify all genotypes that affect susceptibility, and thus shortened the treatment guidance to less than three hours. Additional reagents and reaction time are both needed for subspecies identification and detection of antibiotic resistance when using genetic tests. In contrast, no additional cost or reaction time is needed when we use ML algorithm to detect MABC subspecies based on already existing MALDI-TOF spectra. High cost-effectiveness would be an advantageous and favorable feature for real-world deployment.

Visualization in two- or three-dimensional plot is a common method for demonstration of classification. t-SNE was used to simplify and visualize the complex MS spectra information to a two-dimensional result (**Figure 4**). t-SNE was an unsupervised, non-linear dimension reduction method to process high dimensional data. No apparent clusters or trends could be visualized on the plot, though the discrimination performance is high (**Table 2**). The discordance seems confusing, however, the phenomenon has been reported in several studies. In the research studying ciprofloxacin resistance of *Klebsiella pneumoniae*, t-SNE plot of MALDI-TOF spectra also showed that resistant strain and susceptible strain were not distinguishable, while AUROC of ML model attained 0.89.^31^ In another large scale research of ESKAPE pathogens, all the t-SNE plots of MALDI-TOF spectra showed no specific pattern for discriminating resistant pathogens from susceptible ones.^32^ Likewise, the predictive performance attained as high as 0.95 for detecting oxacillin resistance for *Staphylococcus aureus*. Results of t-SNE and predictive performance of ML models seem inconsistent, but actually the ML algorithms have been proven workable in detecting antibiotic resistance from MALDI-TOF spectrum for many different pathogens. The possible explanation is that the difference of MALDI-TOF spectra between different phenotypes (e.g. drug resistance, subspecies) are subtle. The subtle and implicit difference cannot be detected in low dimensionality (e.g., one, two, or even three dimension). Thus, using ML algorithms for high-dimensional pattern recognition is a reasonable solution.

We used five ML algorithms to construct prediction models. Among the ML methods, DT, RF, and SVM reached over 0.80 in specificity, specificity, accuracy, and AUROC. Not surprisingly, RF outperformed all other ML methods, with highest sensitivity (87%), specificity (87%), accuracy (87%) and AUROC (0.92). The reason for the outperformance could be attributed to its specific bagging method,^33^ which allows the algorithm handle high dimensional data simultaneously without excessive feature selection, as previous studies showed.^19^ Another concern of ML method was that, Wang et al.^34^ mentioned using not independent dataset in model training and validating, could cause issue of overfitting. Nonetheless, in present study, nested cross-validation, which feature selection and model adjustment were conduct with literally different datasets, was used to prevent such problem. On the other hand, LR method revealed the lowest performance in every aspect of model evaluation, with accuracy of 49% and AUROC of 0.5812 (95% CI, 0.5732, 0.5893). This phenomenon demonstrates that conventional statistical algorithm would be not suitable to cope with complex nonlinear MALDI-TOF MS data. ML algorithms could substantially help scientists to analyze data in an efficient and accurate way. Also, well utilization of ML methods in clinical laboratory could save costs and labor.

Teng et al.^13^ proposed that *m/z* 4386.24, 7669.20, 8771.73 were observed in *M. abscessus* subsp. *massiliense*; while *m/z* 7639.70, 8783.84, 9477.48 were observed in *M. abscessus* subsp. *abscessus*. By using cluster analysis, the prediction of subspecies with MALDI-TOF data, the accuracy could reach 100%. Interestingly, they discovered peak of *m/z* 9,477 was characteristic peaks and 100% specific for subsp. *abscessus*, which showed higher resistance to clarithromycin. On the other hand, another research revealed that peak of *m/z* 9,473.31 presented in only 17% among the subsp. *abscessus* isolates and was absent from subsp. *massiliense*.^35^ The peak is suspected to be in accordance with the peak of *m/z* 9,478 that was found in our study (**Figure 6B**). The subtle difference noted between *m/z* 9,473 and 9,478 would be resulted from shifting problem that caused by isotopes, so *m/z* 9,473 and 9,478 could be regarded as the same peptide molecules.^17,20,36^ Base on previous studies, peak of *m/z* 9,478 occurred less than 40% in either subsp. *abscessus* or *subsp. massiliense*.^13^ The low occurring frequency was also noted in our results, indicating that single peak would not be a good feature to distinguish between two subspecies. On the other hand, our findings further highlighted the strength of ML method, taking multiple peak patterns into consideration, and become a more complicated but accurate classifier. Along with other peaks, as shown in **Figure 6B** and previous section, we discovered many essential peaks that were not published by predecessors. Further studies of potential peaks and its corresponding molecules are needed to disclose the full secret of antibiotics resistance of MABC.

Several limitations were found in the present study. One of our limitations was that due to various geographical diversity of microorganism, our single-center study could not generalize directly worldwide. However, by using the proposed ML models could lead to more precise diagnostic result, but also accurate macrolide susceptibility way before the conventional AST results come out. Secondly, among plenty of works, those found discriminating MALDI-TOF MS peaks regarding MABC were distinct; and despite similar protocols and laboratory approach, the results were not easily reproducible. The reason could be the unique biogeographic MS profiles of MABC in different areas, thus eventually caused inconsistent m/z ratio.^33^ We did no aim to propose a perfect research that fit the MABC species data worldwide; on the other hand, we proposed a method that can apply to local epidemiologic and different microorganic data, and thus clinicians can treat the patient according to the results done by local laboratory respectively. Lastly, molecular validations (i.e., protein analysis) data were still lacking, thus could not confirm representing peptides of highlighted m/z ratio peaks in our study.

In conclusion, we presented a method based on MALDI-TOF and ML technologies for rapid and accurate differentiation of macrolide-resistant MABC subspecies from macrolide-susceptible subspecies. The rapid and highly available diagnostic information would guide more precise and adequate treatment of MABC.

## Acknowledgments

This work was supported by Chang Gung Memorial Hospital (CMRPG3M0851, CMRPG3L1011), and the Ministry of Science and Technology, Taiwan (110-2314-B-182A-147 and 111-2320-B-182A-002-MY2).

